# Towards the development of a mucosal vectored vaccine against enterovirus D68

**DOI:** 10.64898/2025.12.12.693937

**Authors:** Anna Zimina, Ekaterina G. Viktorova, Diana Kouiavskaia, Konstantin Chumakov, Amy B. Rosenfeld, George A. Belov

## Abstract

Enterovirus D68 (EVD68) is an emerging pathogen associated with neurological complications, including temporary and permanent paralysis, and lethality. Currently, no EVD68-specific vaccines or antivirals are available. Here, we used a Newcastle disease virus (NDV) vector to develop a live mucosal vaccine that produces EVD68 virus-like particles (VLPs) and evaluated its immunogenicity in a murine model. We made constructs producing VLPs of the prototype EVD68 strain Fermon and the USA/2018-23088 (Ohio-2018) strain from a recent outbreak in the USA. Our results demonstrate robust production and proper processing of EVD68 capsid proteins upon co-expression with the viral protease 3CD. Intranasally-immunized animals effectively seroconverted to NDV antigens, indicating successful replication of the vectors. Antibodies that could bind both the Fermon and Ohio-2018 virions were observed in the sera of mice immunized with either viral vector. At least some animals also demonstrated a strong respiratory mucosal IgA response against EVD68 capsid proteins. These data confirm the immunogenicity of EVD68 proteins expressed from a viral vector in the respiratory tract. Yet neither humoral nor mucosal antibodies were protective against EVD68 infection in cell culture. Analysis of cells infected with recombinant NDVs by electron microscopy indicated structural differences between *bona fide* EVD68 virions and VLPs, suggesting that the lack of neutralizing antibody response is likely due to antigenic disparity between the EVD68 virions and empty VLPs and/or limited stability of VLPs within the murine respiratory tract. These results have important implications for the development of VLP-based vaccines against enteroviruses.

## Introduction

Enterovirus D68 (EVD68), currently classified as a serotype within *Enterovirus deconjuncti* species, was first identified in 1962 in four children in California hospitalized with respiratory distress (Schieble et al., 1967). It remained an obscure pathogen of low interest until 2014, when an outbreak of more than 2000 cases of confirmed EVD68 infections associated with severe respiratory symptoms, mostly in the US and Canada but also in other countries, was reported (Midgley et al., 2015; Poelman et al., 2015; Skowronski et al., 2015). Even more troublesome, some children with EVD68 infections developed neurological symptoms similar to acute flaccid paralysis, the disease previously predominantly associated with poliovirus infections (Greninger et al., 2015). Since 2014, multiple cases of EVD68-related respiratory distress have been registered regularly around the world with biannual peaks, until this pattern was broken in 2020, likely due to social contact restrictions imposed during the SARS-CoV-2 pandemic. The biennial occurrence of EVD68 seems to reemerge in 2022-2024 seasons, however, the number of cases detected was significantly lower than in the pre-SARS-CoV-2 pandemic years (Nguyen-Tran et al., 2025). The spatio-temporal association of EVD68 infections with clusters of non-polio acute flaccid paralysis cases strongly suggests the cause-and-effect relationships (Ayers et al., 2019; Kaye, 2019; Sejvar et al., 2016), although the virus was directly isolated from cerebrospinal fluid from only two patients (Khetsuriani et al., 2006; Kreuter et al., 2011), and only one autopsy report confirms the presence of the viral RNA and proteins in motoneurons (Vogt et al., 2022). The growing concerns over the EVD68 infections, already dubbed “the new polio”, call for the development of countermeasures, including protective vaccines (Cassidy et al., 2018; Helfferich et al., 2019; Morens et al., 2019).

EVD68 replicates in the nasopharyngeal cavity and the upper respiratory tract, similar to rhinoviruses (Blomqvist et al., 2002; Oberste et al., 2004). The seroprevalence and case distribution data indicate global circulation of EVD68 strains, which are currently classified into three major clades (A, B, and C), established according to their divergence from the prototype Fermon strain isolated in 1962, with A and B clades further subdivided into three subclades each (Tokarz et al., 2012). The different clades and subclades demonstrate specific geographical distribution and association with infection of different age groups (Hadfield et al., 2018; Hodcroft et al., 2022). Viruses from subclade B1 were associated with the largest outbreak of EVD68 infections in 2014 in the US (Eshaghi et al., 2017; Greninger et al., 2015; Tokarz et al., 2012). An important question is whether the global diversification of EVD68 strains may eventually result in an antigenic diversity sufficient for establishing different serotypes of the virus. The currently available data are consistent with the existence of one EVD68 serotype, so that the neutralizing antibodies developed against one strain can cross-neutralize others, albeit with different efficacy (Harrison et al., 2019; Karelehto et al., 2019; Patel et al., 2016; Zhang et al., 2018b; Zhang et al., 2015), suggesting that a monovalent vaccine can be broadly protective. EVD68s from different clades replicate similarly in human and murine neurons and astrocytes in cell culture and *in situ* in a brain slice model, although strain-specific differences have been reported (Brown et al., 2018; Rosenfeld et al., 2019).

While no immunocompetent animal model recapitulating the neurological symptoms of EVD68 infection has been established yet, the virus demonstrates neurotropism and induces paralysis in newborn or interferon signaling-deficient mice (Rosenfeld et al., 2019; Sun et al., 2019; Zhang et al., 2018b).

Similar to other enteroviruses, the non-enveloped icosahedral virion of EVD68 contains a single-stranded (+)RNA molecule of ∼7.4 kb coding for one open reading frame. The polyprotein is processed co- and post-translationally by viral proteases into about a dozen final proteolysis products and stable processing intermediates that support the RNA replication and encapsidation of progeny genomes. The precursor of capsid proteins P1 is processed into VP0, VP3, and VP1 proteins, which assemble into pentamer intermediates, followed by the spontaneous formation of stable empty virus-like particles (VLPs) consisting of 60 copies of each of the capsid proteins. RNA encapsidation triggers the final capsid maturation event, the autocatalytic cleavage of VP0 into VP4 and VP2 proteins (reviewed in (Putnak and Phillips, 1981b)).

In this work, we used a vaccine strain of Newcastle Disease virus (NDV) to design a vector expressing the EVD68 capsid protein precursor P1 together with a protease required for its processing, with the idea of generating EVD68 VLPs upon NDV vector replication in the cells of a vaccine recipient. We previously successfully used a similar design for an experimental vaccine against poliovirus, another enterovirus (Viktorova et al., 2018).

NDV is a single-stranded (-)RNA virus. It is an avian pathogen, and the avirulent LaSota strain is used as a live vaccine for poultry throughout the world. NDV is a promising vaccine vector used for the development of several prototypes of human and veterinary vaccines (reviewed in (de Swart and Belov, 2023; Kim and Samal, 2016b)). Importantly, NDV has been extensively investigated as an oncolytic agent, and an NDV-vectored vaccine against SARS-CoV-2 was evaluated in large-scale clinical trials. NDV proved to be safe even in patients with a severely compromised immune system (Freeman et al., 2006; López-Macías et al., 2024; Lundstrom, 2018; Ponce-de-Leon et al., 2023; Schirrmacher, 2016).

The modular genome organization of NDV is highly amenable to the introduction of foreign genes, and the pleomorphic virions of NDV do not have a strict limit for the length of packaged RNA. The replication of NDV in mammalian cells induces potent activation of innate immune signaling because the viral antagonist of interferon response is effective in avian but not in mammalian cells (Park et al., 2003; Vigil et al., 2007). The strong stimulation of innate immune response serves as a built-in adjuvant in NDV-based vaccines, promoting the development of humoral and mucosal immune responses (Kim and Samal, 2016a). NDV infects via the intranasal route and replicates in cells of respiratory mucosal epithelium, which makes it particularly attractive for the development of vaccines against respiratory pathogens such as EVD68.

We designed two NDV vectors expressing capsid proteins of distant EVD68 strains: the historic Fermon strain and the recently isolated USA/2018-23088 (Ohio-2018) strain, associated with a US disease outbreak. Both vectors showed similar replication levels and were sufficiently stable over multiple passages in embryonated chicken eggs. They robustly produced EVD68 capsid proteins, which were properly processed and assembled into VLPs in infected cells. Our data confirms that protease 3CD rather than 3C is required for the processing of enterovirus capsid proteins. No adverse effects were observed upon intranasal administration of recombinant NDVs to C57BL/6J mice. All vaccinated animals developed a strong humoral antibody response to NDV, indicating vector replication. EVD68-binding IgG antibodies were detected in the sera of all immunized mice, and an IgA response against EVD68 capsid antigens was observed in the bronchioalveolar lavage (BAL) of at least some animals, showing that intranasal immunization of mice with NDV-EVD68 can elicit both mucosal and humoral immunity. However, neither serum nor BAL antibodies were able to protect cells in culture from virus infection. Electron microscopic analysis of cells infected with recombinant NDVs indicated structural differences between the EVD68 virions and VLPs. Collectively, our data suggest that the lack of neutralizing antibodies upon vaccination with NDV vectors may be due to structural/antigenic differences between the infectious particle and empty VLPs, and/or VLP instability when produced in the cells of the murine respiratory tract. These results imply that additional structural investigations of EVD68 virions, including the effect of VP0 cleavage, are required for the development of VLPs that more accurately reflect the antigenic properties of the infectious particle.

## Materials and Methods

### Cells and viruses

HeLa and Chicken fibroblast DF-1 cells were grown in Dulbecco’s Modified Eagle Medium (DMEM) supplemented with 10% fetal bovine serum (FBS), 10mM sodium pyruvate, and antibiotic-antimycotic mix (GIBCO), Hep-2 and RD cells were grown in the same medium but without sodium pyruvate.

EVD68 viruses Fermon and USA/2018-23088 (Ohio-2018) were from the laboratory stocks of Dr. Rosenfeld. For propagation, EVD68 was added to RD cell monolayers grown on T-25 or T-75 flasks and allowed to adsorb at 33 °C for 1 hour with gentle rocking. Following adsorption, the cells were incubated in the standard growth medium at 33 °C for about 72 hours.

EVD68 titer was determined by a standard plaque assay as PFU/ml, or by a TCID_50_ method using RD cells grown in 96 wells and calculated using Karber’s formula (Kärber, 1931). For the plaque assay, RD cell monolayers in 6-well plates were washed twice with PBS. EVD68 stock was serially diluted in PBS (prepared in duplicate), and 100 µL of each dilution was added to the appropriate wells. Mock-infected wells received PBS alone. The inoculum was distributed by gently rocking the plates, followed by a 1-hour adsorption at 33 °C. After adsorption, the inoculum was aspirated, and 4 mL of an overlay containing equal parts of 2× DMEM supplemented with 5% FBS and antibiotic-antimycotic, and 1.8% Noble agar (Difco) prepared in molecular grade water was added to each well. Plates were incubated at 33 °C for 5 days before staining with crystal violet.

### Rescue and propagation of recombinant NDVs

Recombinant Newcastle disease viruses were generated using the reverse genetics system for the avirulent strain LaSota described in (Huang et al., 2001). Sequences corresponding to the EVD68 capsid region P1, proteases 3C and 3CD were amplified by PCR, and cloned into the LaSota genome cDNA between P and M, and HN and L genes, respectively. The inserted EVD68 sequences were flanked by NDV transcription start and stop sequences. Cloning details are available upon request. For virus rescue, the plasmid coding for the modified LaSota genome was co-transfected with plasmids coding for NDV N, P, and L genes into HEp-2 cells using Mirus-2020 (Mirus Bio) transfection reagent, and the cells were infected with T7 polymerase-expressing vaccinia virus (Wyatt et al., 1995). After 5 h incubation at 37 °C, the cells were washed once with serum-free DMEM and incubated for 48 hours in DMEM supplemented with 10% allantoic fluid collected from 10-day-old embryonated chicken eggs (Charles River, Manassas, VA), and 2% FBS. The cells were then frozen and thawed once, followed by the clearing of supernatant by low-speed centrifugation. The clarified supernatant was inoculated into the allantoic cavity of 10-day-old embryonated chicken eggs and incubated for 72 hours, after which the allantoic fluid was collected and tested for the NDV HN hemagglutination activity (HA) using chicken red blood cells. HA-positive samples were passaged three more times in embryonated eggs and analyzed for the presence of EVD68 inserts in the NDV RNA by RT-PCR. The NDV expressing EGFP and mCherry proteins was similarly generated previously in our lab.

For isolation of individual NDV clones expressing EVD68 inserts, serially diluted samples of allantoic fluid from infected embryonated eggs were adsorbed for 1 hour at 37 °C on DF-1 cell monolayer grown in 96-well plates, after which DMEM containing 2% FBS and 10% allantoic fluid from non-infected embryonated eggs was added to each well to a final volume of 100 µL. Plates were incubated at 37 °C for 2-3 weeks, with fresh medium added in the middle of incubation, and wells exhibiting cytopathic effect were marked before freezing at −80 °C. After thawing, the contents of each CPE-positive well were diluted 1:4 in PBS, and 150 μL of the dilution was inoculated into the allantoic cavity of embryonated chicken eggs. Allantoic fluid was collected 72 hours post-inoculation, RNA was extracted, reverse-transcribed, and PCR products covering complete EVD68 inserts were sequenced by Sanger sequencing.

For recombinant NDV preparations used for vaccination, allantoic fluid from emdryonated chicken eggs was collected on the third day post-infection, clarified by low-speed centrifugation, and the virus was concentrated by ultracentrifugation at 48,000 × g for 4 hours at 4 °C in a Beckman SW-28 rotor. Pelleted material was resuspended in HBSS containing 2% FBS using a Dounce homogenizer and clarified by sequential low-speed centrifugations (6,000 × g followed by 17,000 × g) to remove large debris. The clarified supernatant was layered onto a pre-formed discontinuous 20-65% sucrose gradient in TNE buffer (100mM NaCl, 10 mM Tris-HCL, 1 mM EDTA) and centrifuged for 2 hours at 24,000 rpm in the SW28 rotor. The visible virus band at the sucrose interface was collected, and the purified virus was diluted 1:1 with HBSS + 2% FBS, aliquoted, and stored at −80 °C. NDV titers were determined as focus-forming units (FFU) by infecting HeLa cell monolayers overnight with serially diluted virus stocks, followed by fixation with 4% paraformaldehyde for 20 min and immunostaining with antibodies against the NDV HN protein. Fluorescent foci were quantified at 40× magnification, using the average of ten fields per well. The titer in FFU/ml was calculated based on the field-of-view area (0.26 mm²).

### Mice immunization

All animal experiments were performed according to the protocols approved by the White Oak consolidated animal program at the FDA. Three-week-old C57BL/6J female mice were obtained from Jackson Laboratory (Bar Harbor, Maine) and were allowed to acclimate for one week. The four-week-old mice were immunized intranasally with 1E6 FFU per mouse of the corresponding vaccine candidates diluted in 30µl of PBS, 15 µL per nostril. Blood samples were collected from the submandibular vein, terminal bleed was performed by cardiac puncture. Bronchioalveolar lavage (BAL) was performed upon euthanasia by washing the respiratory tract with 1 ml of PBS.

### EVD68 Microneutralization assay

The microneutralization assay was performed essentially as described in (WHO, 1997) for poliovirus with minor modifications related to a longer incubation required for EVD68. Briefly, 10 TCID_50_ of EVD68 were preincubated with serial two-fold dilutions of sera in DMEM supplemented with 2% FBS for 3 h, applied to RD cells seeded on 96-well plates, and the medium volume was adjusted to 100µl of DMEM with 10% FBS. Samples were prepared in triplicate. The EVD68 inoculum titer was confirmed in back-titration. Negative control wells were incubated with 100μL DMEM and 10% FBS alone. Plates were incubated at 33 °C for ∼5 days, until the cytopathic effect was observed in the back titration control wells. The cells were then stained with crystal violet.

### ELISA binding antibody assay

Serum IgG antibodies against EVD68 were detected in a Sandwich ELISA assay. High-binding 96-well plates were coated overnight at 4 °C with rat anti-EVD68 (Fermon or Ohio-2018) capture antibodies prepared previously in Dr. Rosenfeld’s lab, diluted in carbonate-bicarbonate buffer, pH 9.5 (SigmaAldrich). Plates were washed four times with TBS-T (50 mM Tris-Cl, pH 7.6; 150 mM NaCl, 0.1% Tween-20), 15 min per wash, and blocked for 4 h with 8% non-fat-dry milk powder (BioWORLD) in TBS-T. EVD68 virions (1E4 PFU) were added to each well and incubated overnight at 4 °C. Following four additional washes with TBS-T and a second 4-hour block, triplicates of serial 10x sera dilutions were added. The next day, plates were washed with TBS-T and incubated with 1:2000 dilution in the blocking solution of HRP-conjugated anti-mouse IgG secondary antibodies (ThermoFisher) for 90 min at room temperature. After four final washes with TBS-T, 100 µL/well of SureBLueTMB HRP substrate (SeraCare KPL) was added. The reaction was stopped after 30 min with 50 µL of 0.2N sulfuric acid, and absorbance was measured at 450 nm.

For the detection of the NDV-binding antibodies, an indirect ELISA was performed. High-binding 96-well plates were coated with 2 µg per well of purified LaSota virions diluted in carbonate-bicarbonate buffer, pH 9.5, and incubated overnight at 4 °C. The next day, plates were washed four times with TBS-T and blocked for 2 hours at room temperature. Sera were added at 2x serial dilutions, each in duplicate, and incubated overnight at 4 °C. Plates were then washed four times with TBS-T and incubated with 1:2000 dilution in blocking solution of HRP-conjugated anti-mouse IgG secondary antibodies for 1 hour at room temperature with low-speed rotation on a shaker platform. The signal detection was performed as described for the Sandwich ELISA assay.

### Mucosal antibody assay

BAL samples were analyzed for the presence of mucosal anti-EVD68 IgA antibodies by Western Blot. HeLa cells were infected with an MOI of 50 of EVD68 Fermon or Ohio-2018, and lysed after 6 or 8h incubation at 33 °C. The lysates were resolved on a 4-15% gradient TGX gel (Bio-Rad), and transferred to a PVDF membrane. Individual membranes containing paired lanes with lysates from EVD68-infected and mock-infected cells were prepared for each BAL sample. The membranes were incubated for 3-4h at 4 °C with BALs diluted 1:1000 in 2% blocking solution in TBS-T buffer, followed by incubation with goat anti-mouse IgA secondary antibody HRP conjugate (Abcam) and developed by ECL Prime Western Blotting Detection Reagent (Cytiva). The images were captured with an Azure Biosystems c300 imager.

### Electron microscopy

Transmission electron microscopy imaging was performed at the University of Maryland School of Medicine’s and School of Dentistry’s Electron Microscopy Core Imaging Facility in Baltimore, Maryland. HeLa cells grown on Thermanox coverslips (EMS) were infected with an MOI of 10 of recombinant NDV expressing EVD68 P1 and 3CD inserts, or NDV expressing EGFP and mCherry proteins (negative control), or an MOI of 50 of EVD68 (positive control), and fixed with 2% paraformaldehyde, 2.5% glutaraldehyde fixative solution in Sodium cacodylate buffer (EMS). Fixative was washed out of the cells with 0.1 M sodium cacodylate buffer. Cells were then post-fixed with 1% osmium tetroxide and 1.5% potassium ferrocyanide in 0.1 M sodium cacodylate buffer. After washing with 0.1 M sodium acetate buffer, cells were *en bloc* stained with 1% uranyl acetate in 0.1 M sodium acetate buffer overnight at 4°C. Then the cells were washed and put through an ascending ethanol series. After washing in propylene oxide, the cells were infiltrated with LX-112 epoxy resin. Coverslips were inverted over resin molds for curing. Thin sections of 90nm were cut from a small blockface placed in an area of dense cells. A transmission electron microscope (JEOL 1400Flash) operated at 80 kV with an AMT CMOS camera was used to image the cells of interest.

### Statistics

Data were analyzed using GraphPad Prism statistical and graphing software. For pairwise sample comparison, an unpaired two-tailed t-test was performed, p-values ≤ 0.05 are designated with *, ≤0.01 with **, ≤0.001 with ***, and ≤0.0001 with ****, NS-non-significant. Graphs show mean value and standard deviation bars.

## Results

### 3CD protease is required for processing of EVD68 capsid protein precursor P1

The genome of NDV is organized in six genes coding for eight proteins (Lamb, 2007). Each gene in the NDV genome is flanked by individual transcription start and end signals, so that the viral polymerase generates separate transcripts for each gene moving from the 3’ end of the (-)RNA genome. This results in a transcription gradient with more transcripts being made for genes located closer to the 3’ end of the genome (Fig. 1A). The precursor of enterovirus capsid proteins P1 needs to be processed into VP0, VP3, and VP1 proteins for virion (or VLP) assembly. In the case of poliovirus, it was shown that the protease 3CD is responsible for the capsid protein processing, rather than the protease 3C, which cleaves most of the polyprotein sites (Ypma-Wong et al., 1988), however, it was not specifically addressed whether 3CD is also important for the capsid protein precursor processing of other enteroviruses. We generated recombinant NDVs with the EVD68 capsid protein precursor P1 inserted between the NDV genes P and M, and the 3CD or 3C proteases between the HN and L genes (Fig. 1A). Due to the NDV transcription gradient, such a design ensures a high level of P1 and a relatively low level of the protease expression, which mitigates cellular toxicity known to be induced by 3C (Barco et al., 2000). These constructs were made with the sequences from the EVD68 strain Fermon. To see if 3CDs can cleave capsids of distantly-related viruses, which may have important implications for enterovirus recombination, another construct was made with the P1 sequence from EVD68 Fermon, and 3CD from poliovirus type I Mahoney (Fig. 1A).

**Figure 1.**
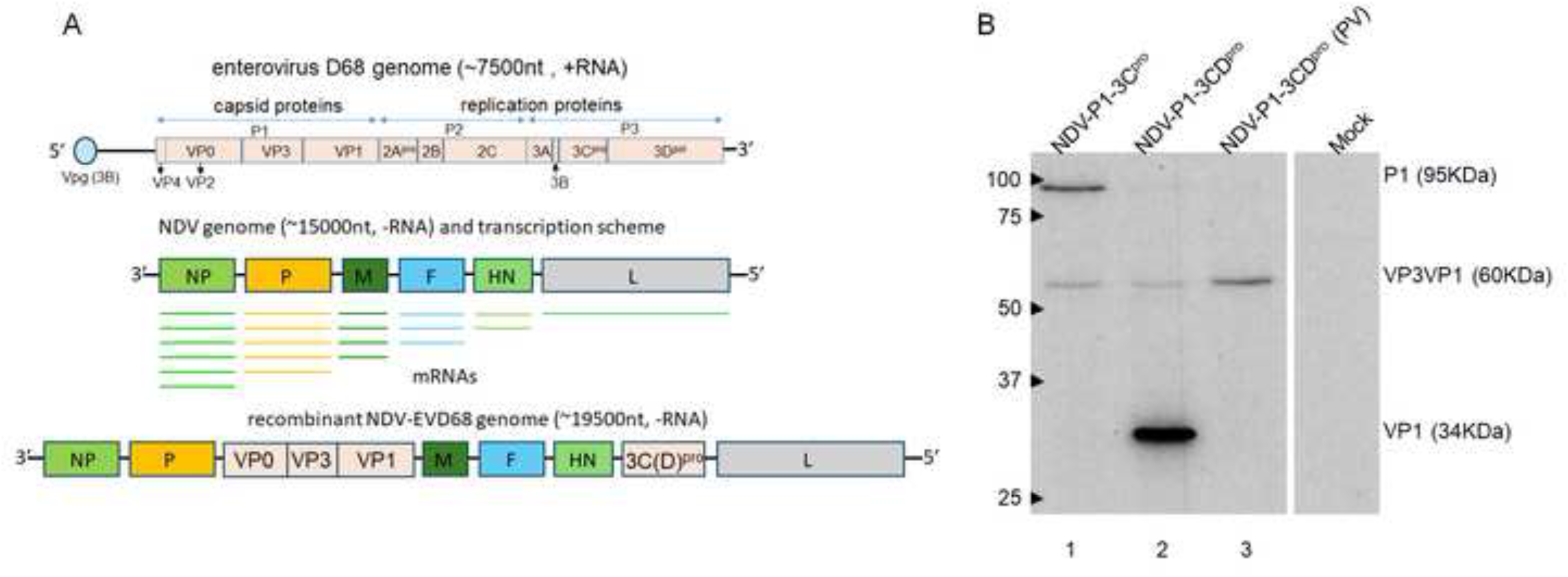
Protease 3CD is required for capsid polyprotein processing of EVD68. **A.** Schemes of the genomes of EVD68 (top, (+)RNA), NDV (middle, (-)RNA), and NDV vector with the inserts of EVD68 precursor of the capsid proteins P1 and proteases 3C or 3CD (bottom, (-)RNA). **B.** Western blot of lysates from HeLa cells infected (and mock-infected control) with NDV vectors expressing EVD68 capsid protein precursor P1 and proteases 3C (1), 3CD (2), or poliovirus protease 3CD (3). The Western blot was developed with an antibody against EVD68 capsid protein VP1. Polyprotein P1 and its processing intermediates, with their molecular weights, are indicated.

All the recombinant NDVs were successfully rescued, and the analysis of P1 expression and processing demonstrated that, like in the case of poliovirus, 3CD protease was the most efficient in the processing of P1 precursor. Interestingly, both EVD68 3C and poliovirus 3CD could partially process P1 at the VP0-VP3 cleavage site, with the polio 3CD protease being even more efficient than the cognate 3C protease of EVD68, as no unprocessed precursor P1 was visible for this construct (Fig. 1B).

Thus, P1 capsid protein processing by the protease 3CD is a conserved feature among enteroviruses. Additionally, this observation suggests that EVD68 and poliovirus 3CDs may similarly cleave other proteins in infected cells.

### The characterization of NDV vectored vaccine candidates against enterovirus D68

To generate a vaccine candidate against a contemporary EVD68 strain, we replaced the P1 sequence in the Fermon-based construct with that of the Ohio-2018 strain, leaving the Fermon 3CD protease. This recombinant NDV was also successfully rescued, and both NDV-Fermon and NDV-Ohio constructs were extensively characterized for EVD68 protein production.

The immunofluorescent analysis of cells infected with the original batch of the recovered recombinant NDVs showed that in the case of NDV-Fermon, only ∼50% of NDV-infected cells produced EVD68 antigen, while in the case of NDV-Ohio, more than 90% of infected cells were positive for EVD68 capsid proteins. We performed a limiting dilution isolation of individual clones from both recombinant NDV pools and characterized them for NDV protein production. One clone for each of the NDV-Fermon and NDV-Ohio that showed close to 100% positivity of EVD68 capsid protein signal was selected for further use as vaccine candidates (Fig. 2A). The EVD68 inserts in the NDV genome were sequenced and confirmed to correspond to the designed constructs. Both constructs effectively produced properly processed capsid proteins of EVD68 (Fig. 2B).

**Figure 2.**
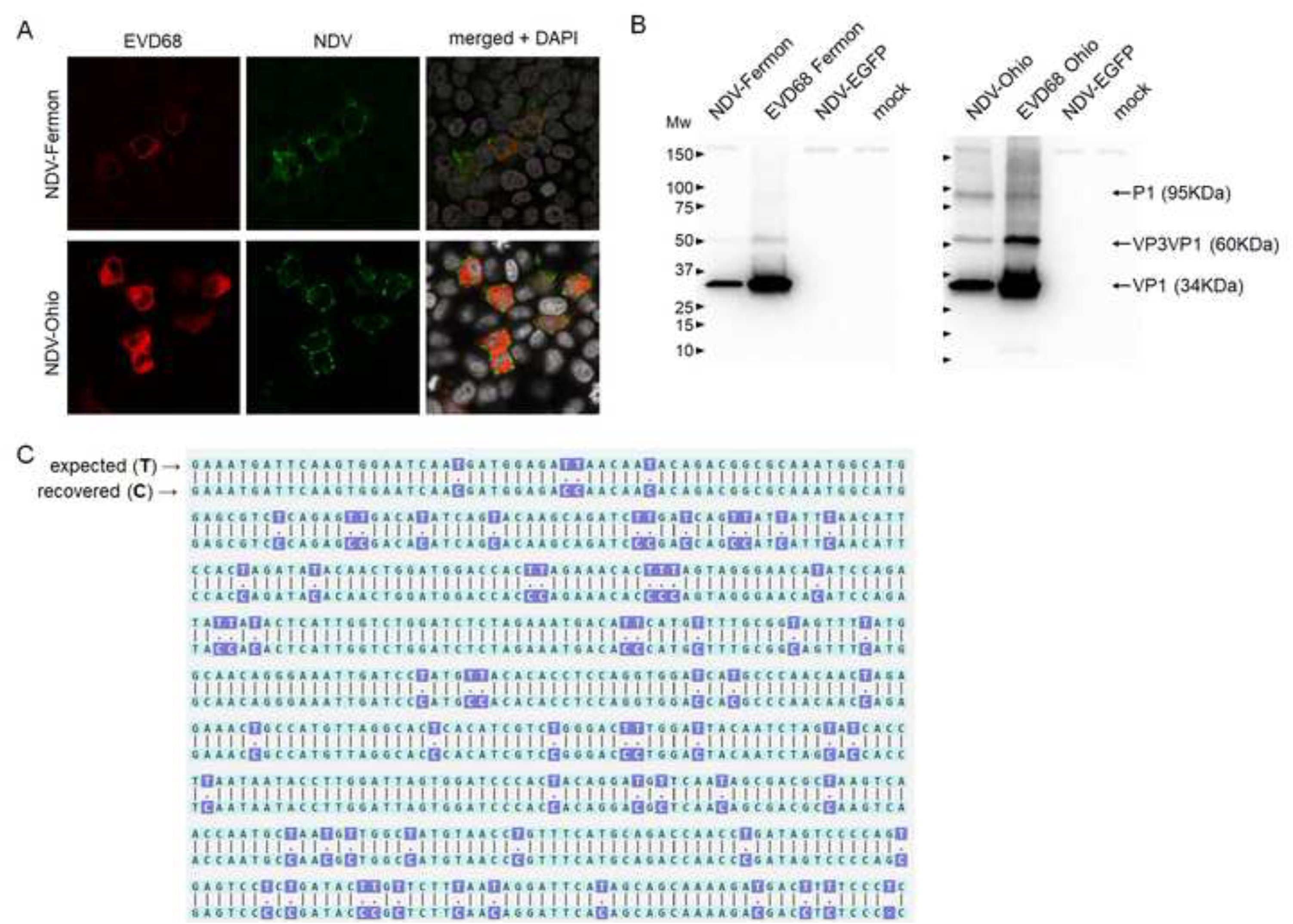
Characterization of NDV vectors expressing capsid proteins of EVD68 strains Fermon and Ohio-2018. **A.** HeLa cells were infected with individual isolates of NDV-Fermon and NDV-Ohio at an MOI of 0.1 and stained with polyclonal antibodies against NDV (green) and a monoclonal antibody against EVD68 capsid protein VP1 (red). **B.** Westen blot of lysates of HeLa cells infected (and mock-infected controls) with NDV-Fermon, NDV-Ohio, control NDV vector NDV-EGFP, and EVD68 strains Fermon or Ohio-2018 developed with an antibody against EVD68 capsid protein VP1. **C.** Comparison of the partial sequence of EVD68 P1 insert in the NDV-Fermon genome between the designed construct (top) and one of the recovered isolates (bottom).

In addition, we also sequenced one NDV-Fermon clone that did not produce EVD68 capsid antigens. Sequence analysis revealed that the isolate had a high frequency of U (T in the cDNA used for sequencing) to C transition mutations, with about 40% of Us in the P1 region changed to Cs (Fig. 2C). This extensive mutagenesis is consistent with the previously described phenomenon of specific silencing of the inserts in the NDV genome attributed to ADAR-1 deamination, an RNA editing process during which adenosines are deaminated to inosine in a double-stranded RNA (de Buhr et al., 2023). This result underscores the importance of the insert sequence verification after the recovery of recombinant NDVs.

### Stability of the recombinant NDV constructs

Stability of a vectored vaccine is essential for its practical applicability. We analyzed the stability of our vaccine candidates after 10 passages in embryonated chicken eggs, a standard system for NDV propagation, which is also broadly used in vaccine manufacturing. Importantly, one passage in the egg comprises multiple replication cycles. Viruses from passages one and 10 were used to infect HeLa cell monolayer at a low MOI so that the expression of NDV and EVD68 antigens could be analyzed in cells infected with individual viral particles (the LaSota strain of NDV cannot spread beyond the originally infected cell in this system without the addition of extracellular proteases). We observed a statistically significant decrease in EVD68 antigen-positive cells upon infection with the NDV vector coding for EVD68 Fermon capsid proteins, from ∼92% to ∼75% for the material from passages one and 10, respectively. The decrease was much less prominent in the case of NDV-Ohio, from ∼94% to ∼85%, which did not reach statistical significance (Fig. 3). These results suggest a sequence-specific selection pressure on recombinant NDVs and are consistent with the previous observation of a much more pronounced editing of the Fermon P1 insert during the recombinant NDV virus recovery.

**Figure 3.**
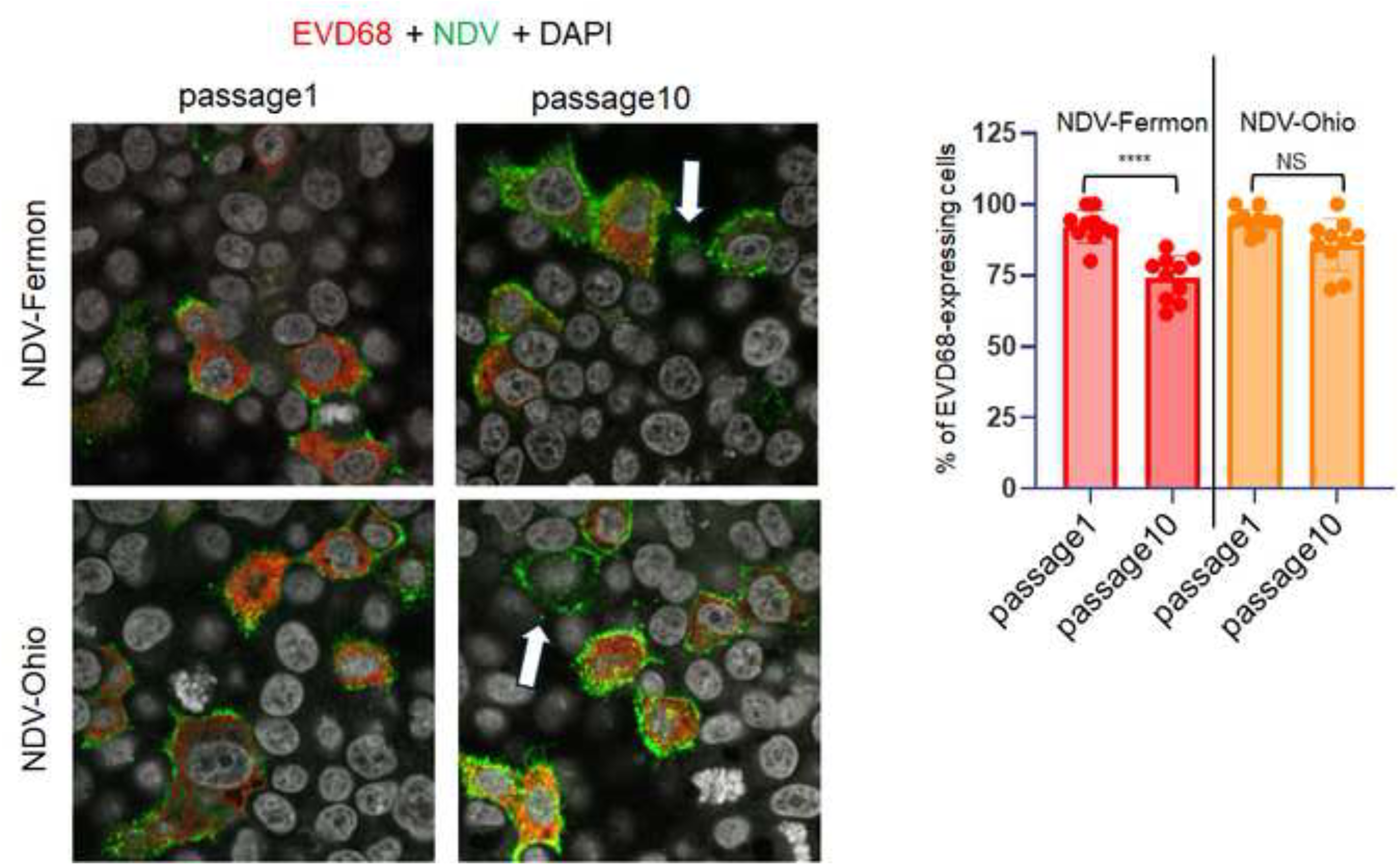
Stability of NDV vectors expressing capsid proteins of EVD68 strains Fermon and Ohio-2018. Left, representative images of HeLa cells infected with a low MOI of NDV-Femon and NDV-Ohio isolates after passage 1 and 10 in embryonated chicken eggs. The cells were stained with polyclonal antibodies against NDV (green) and a monoclonal antibody against EVD68 capsid protein VP1 (red). Arrows indicate cells that express NDV antigens but not those of EVD68 in the material from the 10^th^ passage. Right – quantitation of cells expressing both antigens. Dots represent data for individual images, with ∼200 total number of infected cells.

### Characterization of the humoral immune response upon intranasal delivery of recombinant NDV expressing EVD68 capsid proteins

As no immunocompetent small animal model is available for the assessment of the protective efficacy of anti-EVD68 vaccine candidates, we evaluated the immune response elicited by the recombinant NDVs in adult C57BL/6J mice. The animals were immunized intranasally with 1E6 infectious units of NDV expressing EVD68 capsid proteins, one control group was immunized with an NDV vector with EGFP and dsRed coding genes in place of the EVD68 sequences (NDV-EGFP), and the other control group was sham-immunized with PBS. In total, the vaccine was administered four times, and the blood was collected after each immunization. After the final immunization, bronchoalveolar lavage (BAL) was collected, and the animals were terminally bled (Fig. 4A).

**Figure 4.**
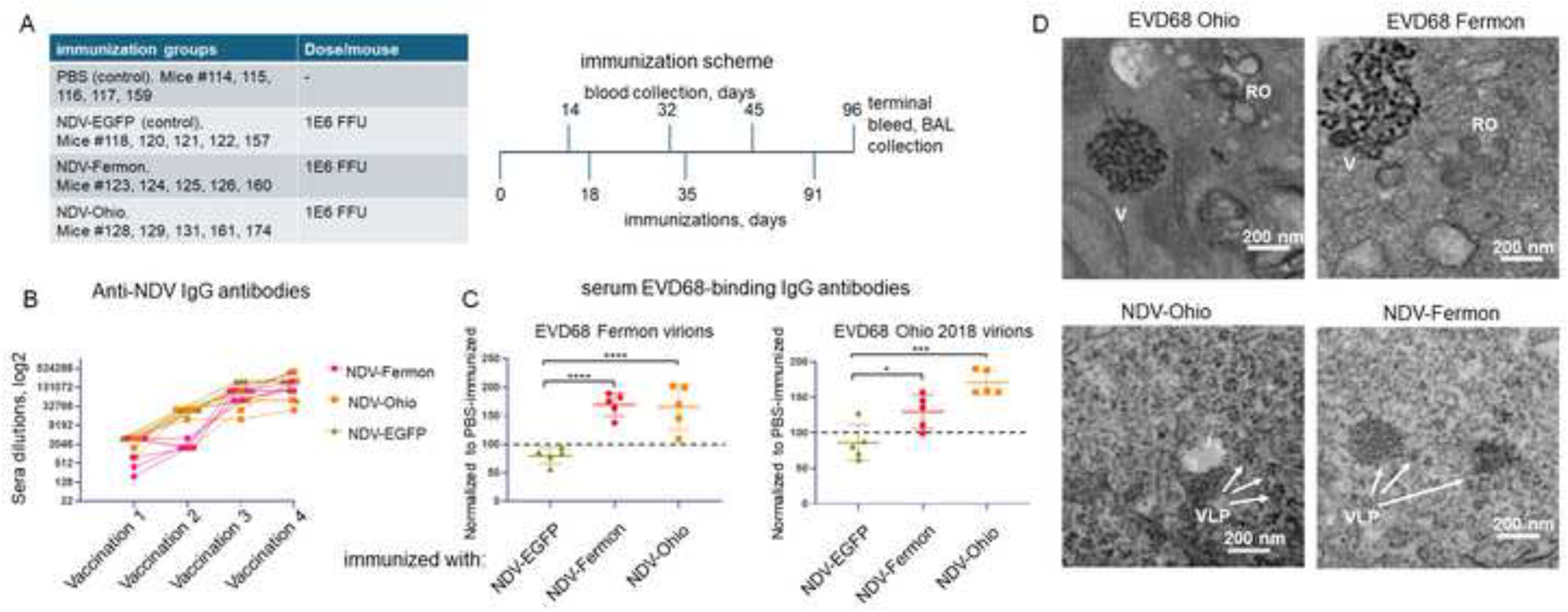
The development of humoral immune response upon intranasal immunization with NDV vectors expressing capsid proteins of EVD68 strains Fermon and Ohio-2018. **A.** Animal groups used in the immunization experiment and immunization scheme. Five C57BL/6J mice per group were immunized with NDV vectors expressing EVD68 capsid proteins, a control NDV-EGFP vector, or sham-immunized with PBS. The viruses were delivered intranasally on days 0, 18, 35, and 91, and blood was collected on days 14, 32, and 45. On day 96 post first immunization, the animals were sacrificed, and the terminal blood and BAL collection was performed. **B.** The development of the immune response against NDV antigens. The serum anti-igG antibody levels were evaluated in an indirect ELISA assay. The Y-axis shows sera dilutions that generated the ELISA signal exceeding the average signal from sera from mice sham-immunized with PBS by more than twice the standard deviation of the control samples. Points on the graph represent individual animals, connecting lines allow tracing the dynamics of individual animal response. **C.** A Sandwich ELISA assay evaluation of antibodies binding to EVD68 virions of Fermon (left) and Ohio-2018 (right) strains in sera collected at the terminal bleed. The signal from sera from mice immunized with the NDV vectors (control NDV-EGFP, NDV-Fermon, and NDV-Ohio) is normalized to the average signal from sera from mice sham-immunized with PBS. Individual animal responses are shown. **D.** Transmission electron microscopy images of cells infected with EVD68 strains Ohio-2018 and Fermon (top), and NDV vectors expressing corresponding capsid proteins (bottom). Note the difference between electron-dense clusters of EVD68 virions (V) located close to replication organelles (RO), and more electron-transparent VLPs.

First, we analyzed the development of the antibody response against the NDV vector. As can be seen from Fig. 4B, all mice that received NDV vectors developed a robust anti-NDV signal in an ELISA assay, which was evident after the first immunization and continued to increase with the subsequent immunizations. However, we could not detect the neutralizing anti-EVD68 antibodies in either sera or BAL collected from the immunized animals. Nevertheless, we detected EVD68 virion-binding antibodies in the sera collected after the fourth immunization (Fig. 4C). Sera from animals immunized with NDV-Fermon and NDV-Ohio vectors contained binding antibodies against EVD68 virions of both strains, indicating their antigenic similarity (Fig. 4C). These data demonstrate that intranasal production of EVD68 capsid proteins is able to elicit a humoral response in C57BL/6J mice.

The absence of detectable neutralizing antibodies may be due to structural differences between the infectious particle and VLP. We analyzed cells infected with EVD68 and those infected with NDV vectors expressing EVD68 capsid proteins by electron microscopy (Fig. 4D). In the former, compact clusters of electron-dense particles were often detected close to the viral replication organelles identified as assemblies of single and double-membrane vesicles in the cytoplasm. In the cells infected with NDV-EVD68, such clusters were rare, and the particles appeared more transparent and scattered in the cytoplasm (Fig. 4D). We interpret these data as evidence of significant structural differences between the *bona fide* EVD68 virions and VLPs produced in this system, although the limitations of standard EM imaging for the analysis of small enterovirus particles should be taken into account.

### Characterization of the mucosal immune IgA response upon intranasal delivery of recombinant NDV expressing EVD68 capsid proteins

To determine if intranasal immunization with NDV vectors induced the development of mucosal immune response, we performed a Western blot-type analysis with the lysates from cells infected with EVD68 (which contain all viral proteins), using BAL as a source of primary antibodies, and developing the signal with an anti-mouse IgA secondary antibody HRP conjugate. No EVD68-specific signal was detected in BALs of animals immunized with either NDV-EGFP or sham-immunized with PBS, as expected (Fig. 5, right panels). At the same time, BALs from three out of five animals immunized with either NDV-Fermon or NDV-Ohio produced highly specific signals with the lysates from EVD68-infected cells compared to those from mock-infected controls (Fig.5, left panels). The bands corresponding to the molecular weights of EVD68 capsid polyprotein precursor P1, processing intermediates VP1VP3 and VP3VP0, as well as fully processed VP1, could be detected. Interestingly, BALs from some animals also contained antibodies reactive with ∼50KDa band in the lysates from EVD68-infected cells, which can indicate the development of 3D-specific antibodies (3D is expressed by NDV vectors as part of 3CD). The BALs from the remaining animals did not produce easily interpretable results. Thus, the intranasally-delivered NDV vectors can induce a mucosal IgA immune response against EVD68 capsid proteins.

**Figure 5.**
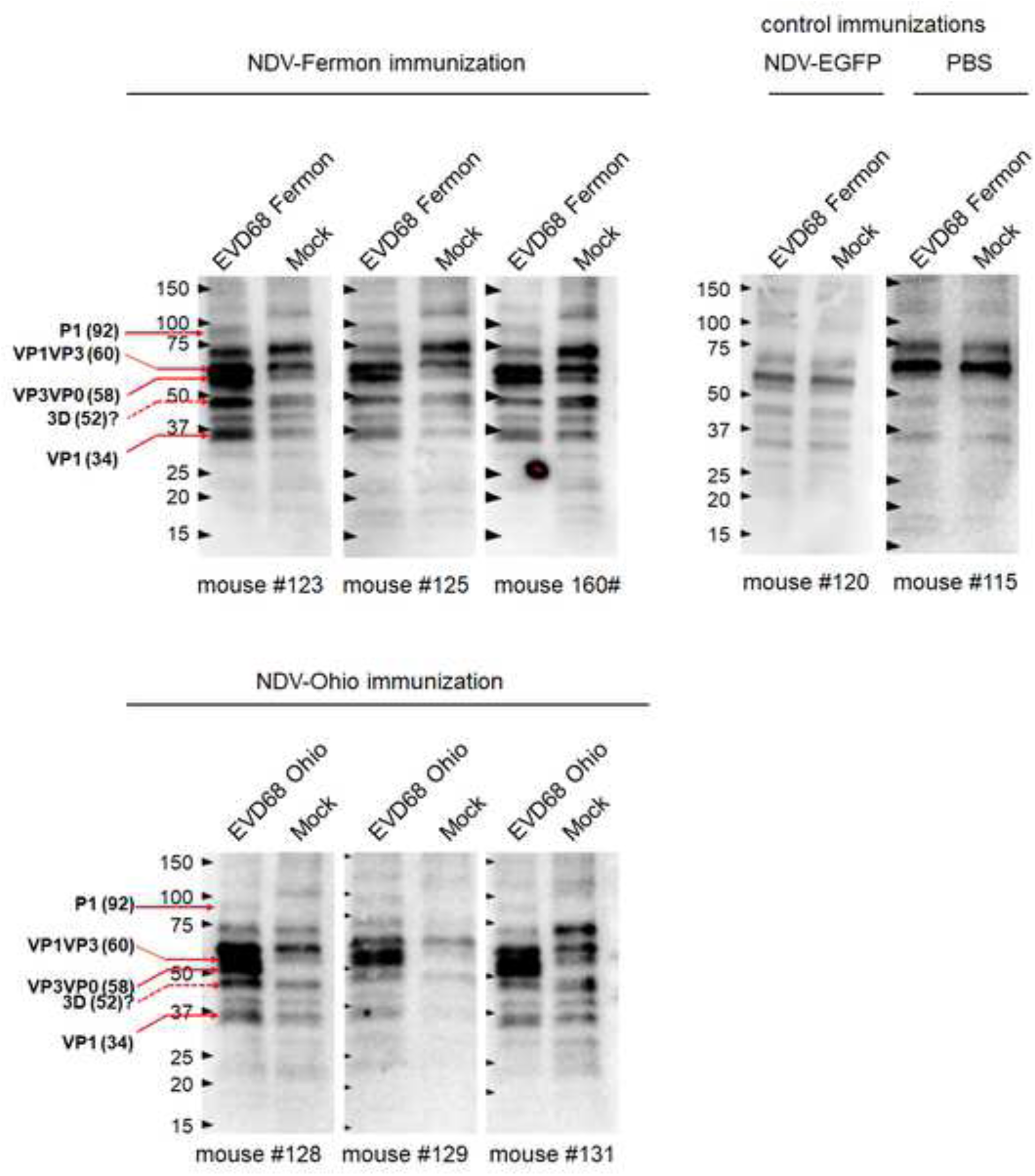
The development of mucosal immune response upon intranasal immunization with NDV vectors expressing capsid proteins of EVD68 strains Fermon and Ohio-2018. Lysates from EVD68-infected (or mock-infected control) HeLa cells were resolved in gradient 4-15% polyacrylamide gels and analysed in a western blot assay with BAL as the source of primary antibodies and anti-mouse IgA secondary antibody HRP conjugate. Note the specific viral protein bands recognized by antibodies from BALs of animals immunized with NDV vectors expressing EVD68 capsid proteins (left panel), but not with the control NDV-EGFP vector or sham-immunized with PBS. For control immunizations, representative images are shown.

## Discussion

Among more than 300 known enteroviruses associated with human diseases, licensed vaccines are currently available only against poliovirus (worldwide) and Enterovirus A71 (in China and some other countries) (Minor, 2014; Reed and Cardosa, 2016). The long history of massive polio vaccination with the first generation of inactivated Salk vaccine (IPV), and live attenuated Sabin vaccine (OPV), licensed in the US in 1955 and 1963, respectively, attests to the efficacy and safety of these vaccines. Recently, improved versions of poliovirus vaccines, such as live attenuated strains with better genetic stability, and the inactivated vaccines derived from attenuated rather than original wt strains have been introduced into clinical practice (Bakker et al., 2011; Ochoge et al., 2024). In principle, such traditional vaccines could be developed against any other enterovirus. Yet, creating genetically stable attenuated strains or devising inactivation methods that preserve protective virion antigens are time-consuming, and the results are not guaranteed, since each new virus requires its own trial-and-error approach. Moreover, even for highly successful anti-poliovirus vaccines, the safety and economic limitations of IPV and OPV have been exceedingly recognized, prompting research into alternative technologies that can rapidly and reliably deliver safe and efficacious vaccines against old and emerging enterovirus threats.

The protective enterovirus antigens often encompass spatial epitopes in the icosahedral virions (Minor et al., 1986; Minor et al., 1983; Page et al., 1988; Vogt et al., 2020), which makes the VLP-based technologies the most attractive direction for the development of new, safer vaccines aimed at inducing neutralizing antibodies. VLPs are normally produced during enterovirus infections (Maizel et al., 1967; Putnak and Phillips, 1981a), and at least for some enteroviruses, it has been shown that VLPs retain the protective antigen conformation of the virions as evidenced by the induction of neutralizing antibodies upon VLP immunization (Hassine et al., 2020; Liu et al., 2012; Zhang et al., 2012). Enterovirus VLPs can spontaneously form upon synthesis of capsid proteins alone, thus allowing their production without the propagation of pathogenic enteroviruses, which is economically attractive for vaccine manufacturing due to reduced biosecurity requirements. Poliovirus VLPs, for example, have been successfully produced in different industrial-level expression systems, such as yeast, plants, and insect cells (Rombaut and Jore, 1997; Sherry et al., 2023; Urakawa et al., 1989). Yet, enterovirus VLPs are intrinsically less stable than virions since the final maturation cleavage of VP0 into VP4 and VP2 is triggered only upon RNA packaging, and RNA inside the virion likely additionally stabilizes the proteinaceous shell (Basavappa et al., 1994; Hummeler et al., 1962). Thus, different approaches to increase the VLP stability are actively being pursued, such as selection of thermostable mutations and protein modeling-predicted stabilization (Adeyemi et al., 2017; Fox et al., 2017; Kingston et al., 2022; Sherry et al., 2025).

Here, we explored another approach to VLP-based vaccination by producing the P1 precursor of capsid proteins of EVD68 together with the protease 3CD required for its processing from an NDV vector. Such a system allows the generation of enterovirus VLPs directly in the cells of a vaccine recipient, which eliminates the requirements of their preliminary production and purification. Moreover, VLPs are presented to the immune system in the context of NDV vector replication, which stimulates innate anti-viral signaling, serving as an intrinsic adjuvant promoting the immune response. Since NDV is an avian-specific virus, pre-existing anti-NDV antibodies are virtually absent in the general population (Charan et al., 1981; Howitt et al., 1948; Pedersden et al., 1990). Moreover, NDV is a member of the genus *Othoavulavirus,* which includes at least 10 other similar avian-restricted virus species, which allows the development of antigenically distinct viral vectors by either using those viruses directly or replacing the NDV glycoproteins with those from other orthoavuloviruses (Bui et al., 2017; Samuel et al., 2011; Tsunekuni et al., 2017). Thus, orthoavulovirus-based vectors are highly amenable for either creating vaccines against different pathogens or using antigenically different vectors for sequential vaccinations with the same protective antigens.

Another important advantage of the NDV vector is that it can be delivered intranasally for replication in the respiratory tract, which is attractive for the development of vaccines against respiratory pathogens like EVD68. Indeed, the efficacy of an NDV-vecored vaccine against influenza viruses has been experimentally demonstrated in mammalian systems (DiNapoli et al., 2010), and the NDV vaccine against SARS-CoV-2 showed promising results in clinical trials (Lopez-Macias et al., 2025).

We evaluated the immunogenicity of our construct upon intranasal delivery in adult mice. The animals seroconverted to NDV antigens, indicating vector replication, but surprisingly, we could not detect neutralizing anti-EVD68 antibodies in either sera or BAL. We only detected humoral EVD68-binding antibodies after the fourth immunization. One possible explanation for the absence of a detectable antibody response following previous immunizations is that EVD68 capsid proteins are less immunogenic than NDV glycoproteins, and/or an insufficient level of NDV infection upon delivery to the murine respiratory tract. The limitations of murine models for the assessment of the efficacy of NDV vectored vaccines have been noticed previously (DiNapoli et al., 2007a; DiNapoli et al., 2007b).

Previous reports indicated that ajuvanted vaccines prepared with purified EVD68 VLPs expressed in yeast, mammalian, or insect cells can induce neutralizing antibodies upon intramuscular or intraperitoneal immunization in mice or non-human primates (Dai et al., 2018; Krug et al., 2023; Zhang et al., 2018a). The difference with our results could likely be attributed to the high doses of VLPs used in these studies, their stabilization upon adsorption on the adjuvant, and different immunization routes. Nevertheless, the detection of EVD68-binding humoral IgGs and mucosal IgAs against EVD68 capsid antigens confirms the mucosal immunogenicity of EVD68 proteins produced by the NDV vectors. In this regard, it is possible that combining the intramuscular prime delivery of the NDV-EVD68, which should increase the efficacy of infection, with the respiratory boost can improve the immunogenicity of these vaccine candidates.

Our data do not allow making conclusions about attributing Fermon and Ohio-2018 strains to the same serotype, but the cross-reactivity of binding antibodies suggests their antigenic similarity. While it is not currently established what level of neutralizing antibodies is required for protection against EVD68-induced diseases, the available reports indicate that immunization with EVD68 VLPs can induce at least some level of neutralizing antibodies against heterologous strains (Krug et al., 2023), which suggests that monovalent vaccines can be broadly protective. Although neutralizing antibodies are considered the best biomarker for protection against the development of severe disease following enterovirus infection, the protective efficacy of non-neutralizing anti-enterovirus antibodies has also been demonstrated (Du et al., 2023), and it would be interesting to determine if mucosal non-neutralizing antibodies could be protective against EVD68.

Our data suggest that the lack of neutralizing antibody response is likely due to structural and, therefore, antigenic differences between VLPs expressed in our system and EVD68 virions. EVD68 is adapted to replication in the human upper respiratory tract, with the optimal temperature of EVD68 replication of 33 °C (Freeman et al., 2021; Oberste et al., 2004). Mice, on average, have a higher body temperature than humans (Abreu-Vieira et al., 2015), which may have destabilized EVD68 VLPs. Interestingly, EVD68 VLPs that successfully induced neutralizing antibodies were produced at lower temperatures in yeast and insect cell expression systems (Dai et al., 2018; Zhang et al., 2018a).

The promising direction for improving the vectored EVD68 vaccine is the incorporation of sequences coding for capsid protein mutants that would generate VLPs stabilized in antigenically correct conformation, similar to the stabilized VLPs developed for experimental poliovirus and enterovirus A71 vaccines (Adeyemi et al., 2017; Basavappa et al., 1994; Fox et al., 2017; Kingston et al., 2022; Sherry et al., 2025).

Overall, these results confirm our previous data that the NDV vector is well-suited for the rapid development of anti-enterovirus vaccines based on *in vivo* expression of VLPs (Viktorova et al., 2018), but also underscore the significant differences in the enterovirus VLP stability and immunogenicity. Such vaccines can be easily delivered via the intranasal route and induce a strong mucosal immune response, which makes them an attractive addition to the extremely limited arsenal of anti-enterovirus measures.

## Acknowledgements

The work was supported by the NIH grant R21AI153976 to GAB. AZ was supported by the Basil & Anne Hatziolos Scholarship Fund for Veterinary Medical Research. EM studies were supported by the Mark Kukucka Fund.

